# How many individuals share a mitochondrial genome?

**DOI:** 10.1101/374686

**Authors:** Mikkel M Andersen, David J Balding

## Abstract

Mitochondrial DNA (mtDNA) is useful to assist with identification of the source of a biological sample, or to confirm matrilineal relatedness. Although the autosomal genome is much larger, mtDNA has an advantage for forensic applications of multiple copy number per cell, allowing better recovery of sequence information from degraded samples. In addition, biological samples such as fingernails, old bones, teeth and hair have mtDNA but little or no autosomal DNA. The relatively low mutation rate of the mitochondrial genome (mitogenome) means that there can be large sets of matrilineal-related individuals sharing a common mitogenome. Here we present the mitolina simulation software that we use to describe the distribution of the number of mitogenomes in a population that match a given mitogenome, and investigate its dependence on population size and growth rate, and on a database count of the mitogenome. Further, we report on the distribution of the number of meioses separating pairs of individuals with matching mitogenome. Our results have important implications for assessing the weight of mtDNA profile evidence in forensic science, but mtDNA analysis has many non-human applications, for example in tracking the source of ivory. Our methods and software can also be used for simulations to validate models of population history in human or non-human populations.

**Author Summary:** The maternally-inherited mitochondrial DNA (mtDNA) represents only a small fraction of the human genome, but mtDNA profiles are important in forensic science, for example when a biological evidence sample is degraded or when maternal relatedness is questioned. For forensic mtDNA analysis, it is important to know how many individuals share a mtDNA profile. We present a simulation model of mtDNA profile evolution, implemented in open-source software, and use it to describe the distribution of the number of individuals with matching mitogenomes, and their matrilineal relatedness. The latter is measured as the number of mother-child pairs in the lineage linking two matching individuals. We also describe how these distributions change when conditioning on a count of the profile in a frequency database.

## Introduction

Human mitochondrial DNA (mtDNA) has long been a useful tool to identify war casualties and victims of mass disasters, the sources of biological samples derived from crime scenes or to confirm matrilineal relatedness [1, 2, 3]. The autosomal genome is much larger and has higher discriminatory power, but the mitochondrial genome (mitogenome) has multiple copies per cell, allowing better recovery of sequence information from degraded samples [1, 3], including ancient DNA [4, 5]. In addition, some biological samples such as fingernails, old bones, teeth and hair have mtDNA but little or heavily degraded autosomal DNA.

It has now become widely feasible to sequence all 16,569 mitogenome sites as part of a forensic investigation [6, 7, 8]. For autosomal short tandem repeat (STR) profiles, there are two alleles per locus and because of the effects of recombination, the alleles at distinct loci are treated as independent, after any adjustments for sample size, coancestry and direct relatedness [9]. In contrast, the maternally-inherited mitogenome is non-recombining, behaving like a single locus at which many alleles, or haplotypes, can arise. Due to finite population size and relatedness, the variation in mitogenomes in any extant population is greatly restricted compared with what is potentially available given the genome length. Whereas a match of two mitogenomes without recent shared ancestry is in effect impossible, there can be large sets of individuals sharing the same mitogenome due to matrilineal relatedness that is distant compared with known relatives but much closer than is typical for pairs of individuals in the population.

This limited variation has important implications for the use of mtDNA to help identify individuals or establish relatedness. A match between the mtDNA obtained from bones found under a Leicester UK carpark and a living matrilineal relative of the former King of England, Richard III, played an important role in establishing the bones as those of the king. However, in contrast with popular reports of genetic evidence “proving” the identification, the mtDNA evidence was not decisive, contributing a likelihood ratio (LR) of 478 towards an overall LR of 6.7 million in favour of the identification [10]. Although that mitogenome was at the time unobserved in the available databases, its observation in both the skeleton and a contemporary individual meant that it was expected to exist in hundreds and perhaps thousands of others. The public interest in the story led to multiple matches being subsequently observed in contemporary individuals, raising the question of how many humans alive today share this “royal” mitogenome?

We recently addressed similar questions for paternally-inherited Y chromosome profiles [11]. Forensic Y profiles focus on a few tens of STR loci, but these can have a combined mutation rate as high as 1 per 7 generations [11, 12], much higher than the mutation rate for the entire mitogenome, for which estimates range up to around 1 per 70 generations (see Materials and Methods). We showed that the high mutation rate of Y profiles has dramatic consequences for evaluating weight of evidence. For example, males with matching Y profiles are related through a lineage of up to a few tens of meioses. Further, the number of males with a matching Y profile varies only weakly with population size, and since the population size relevant to a forensic identification problem is typically unknown, it follows that the concept of a match probability that can be useful for autosomal DNA profiles is of little value for Y profiles.

Because of the lower mutation rate for the mitogenome, the situation is less extreme for mtDNA profiles than for Y profiles. Here we describe the distribution of the number of individuals with the same mitogenome as a randomly-chosen individual under three demographic scenarios and two mitogenome mutation models, finding that the number is typically of the order of hundreds rather than the tens that share a Y profile. The number of mitogenome matches is consequently more sensitive to demographic factors than is the case for Y profiles, but it remains a small fraction of the population relevant to a typical crime scenario. As we did previously for Y profiles, we also describe the conditional distributions given database frequencies for the observed mitogenome, assuming that the database is randomly sampled in the population. We show for example that a mitogenome that is unobserved in a large database can nevertheless exist in hundreds of individuals in the population. We also show that individuals sharing a mitogenome are related, typically within up to a few hundred meioses, which is much more distant than recognised relationships but still much closer than the relatedness of random pairs of individuals in a large population. Therefore the matching individuals may not be well-mixed in the population so that database statistics can be an unreliable guide to the number of matching individuals in the population.

## Results

See Materials and Methods for details of our two mutation models, based on [13] and [14], and three demographic scenarios which we denote 1.2M growth, 1.2M constant and 300K constant.

As for Y profiles, it is difficult to rigorously check our simulation models against empirical databases because real-world databases often result from informal sampling schemes that are far from random samples. They are often drawn from a much larger population than is relevant to a specific crime scenario, and sometimes from a number of different administrative regions such as states. However, broad-brush comparisons are useful and for this purpose we identified a US Caucasian database of 263 mitogenomes [15], which includes 259 distinct haplotypes, a very high level of diversity (259/263 = 98%) that reflects sampling from many US states. All our simulated databases of size 263 show less haplotype diversity than this database, but those under the 1.2M constant model come close (Figs 1 and A1). We also considered an Iranian database [16] of size 352 with 315 distinct haplotypes (89% diversity). This total included several distinct ethnic identities: Persians (181, 91% diversity), Qashqais (112, 84% diversity) and Azeris (22, 100% diversity). The simulated databases of size 352 under the 1.2M growth and 300K constant models show mtDNA diversity close to that of the Iranian database.

**Figure 1:**
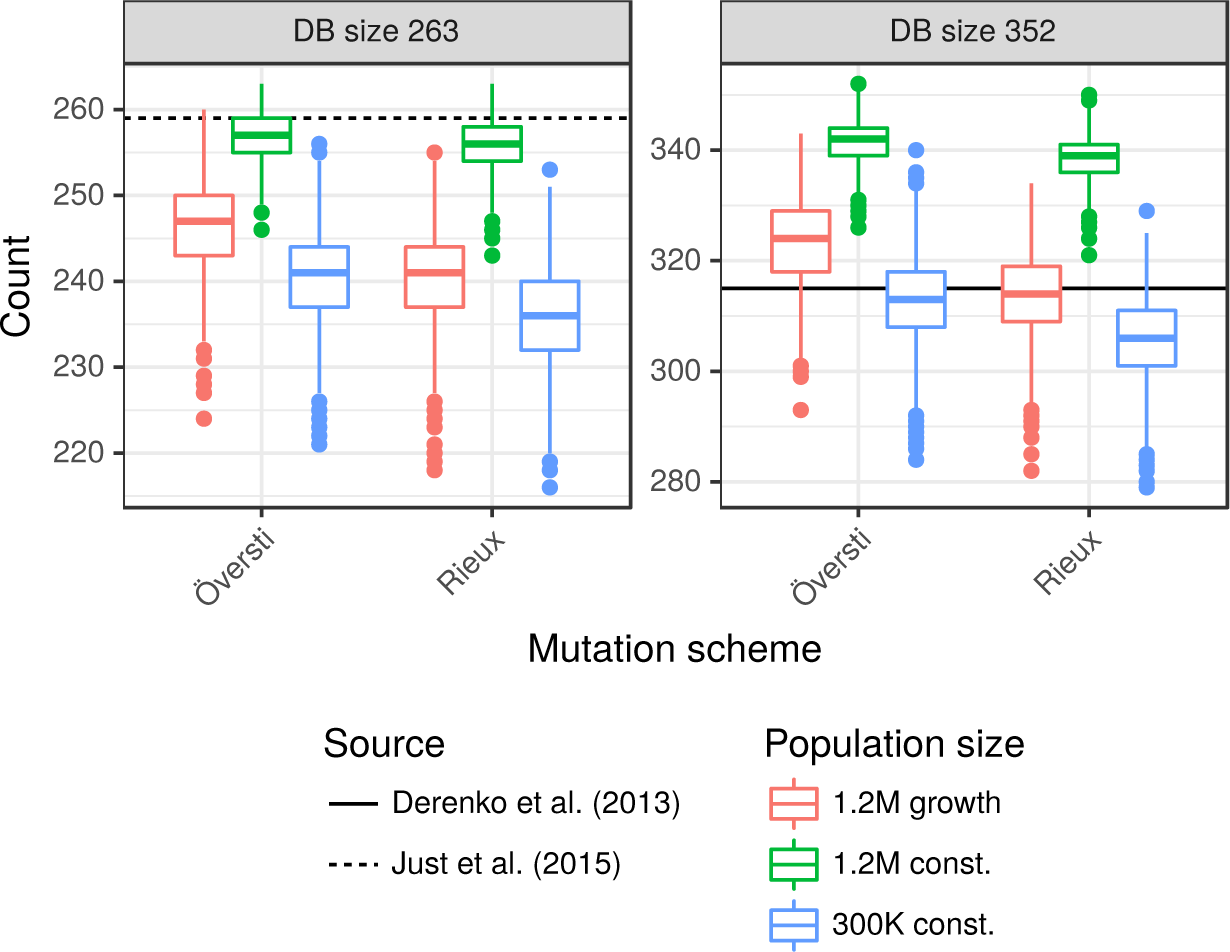
Comparison of simulated with US and Iranian databases. Boxplots show the distribution of the number of distinct haplotypes arising from 2,500 random databases of sizes 263 and 351 obtained under our three demographic and two mutation models. The horizontal reference lines show the numbers of distinct haplotypes in US [15] and Iranian [16] databases of those sizes. See Fig. A1 for distributions of the numbers of singletons and doubletons.

Low mitogenome diversity has been reported in three Philippines ethnic groups with 39, 43 and 27 mitogenomes yielding a diversity of 51%, 58% and 81% [17], which may reflect low population size and isolation. These lower levels of diversity may be appropriate in some forensic contexts, and would require different demographic models from those presented here.

For both mutation schemes, Fig. 2 (black curves, which are the same in each row) shows the cumulative distribution of the number of mitogenomes in the live population matching that of the PoI (person of interest). The distributions (see Table 1 for quantiles) are similar for the 1.2M and 300K constant models (middle and right columns), with the number of sequence matches with the PoI almost always < 1,000, but for 1.2M growth model some PoI have > 5,000 matches.

**Figure 2.**
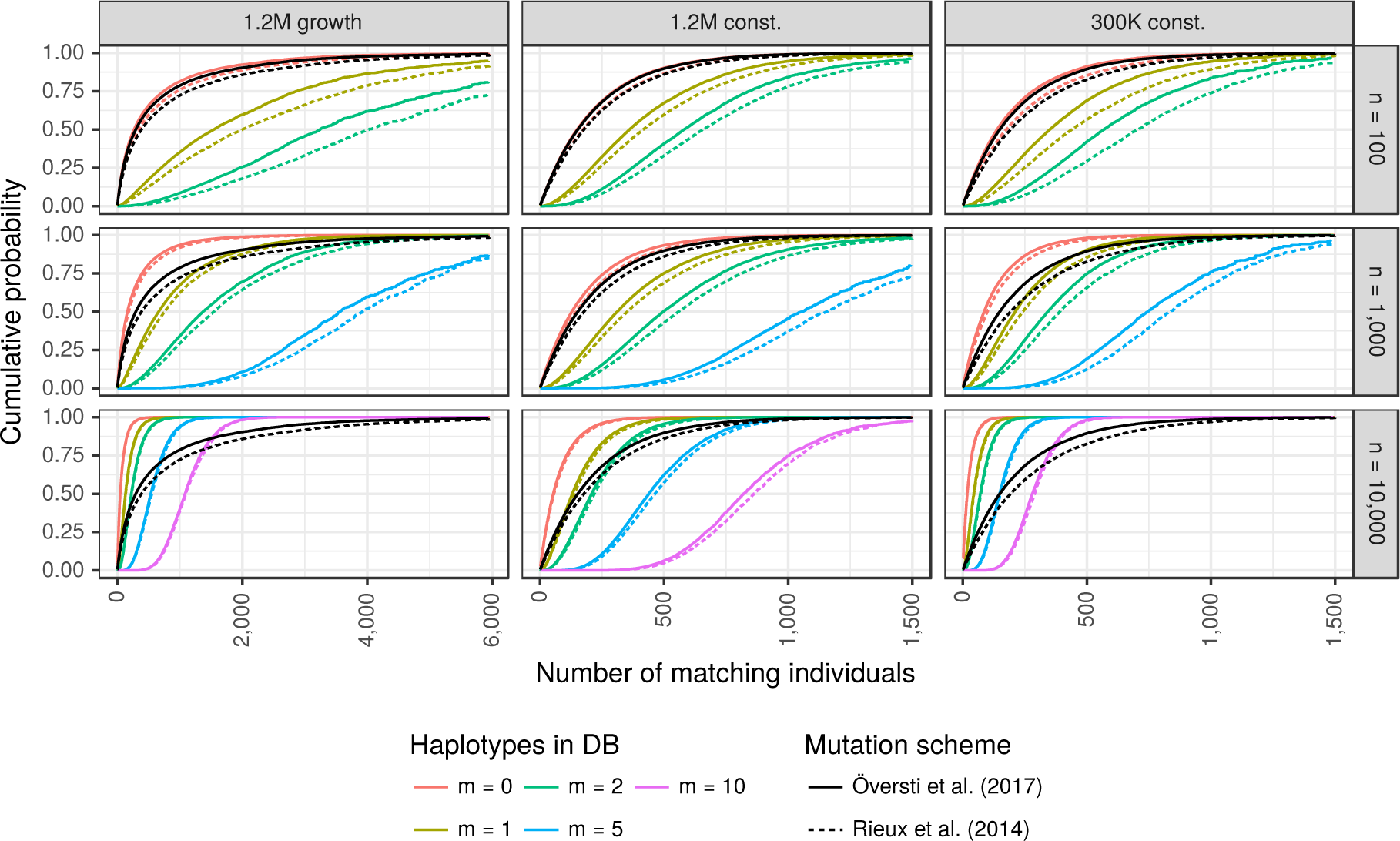
Cumulative distributions of the number of matching individuals. Black lines show unconditional distributions. Coloured lines show the distributions conditional on *m* matching mitogenomes in a reference database of size *n*, for up to five values of *m* (see legend for colour codes) and three values of *n* (one per row). Quantiles of the distributions shown in the middle column are given in Tables 2 and A3 for the mutation models of [13] and [14], respectively. See text for references to additional tables for the other demographic scenarios.

**Table 1.** Estimated quantiles of the number of matching individuals. Key quantiles of the unconditional distributions (black curves of Fig. 2).

| Demographic scenario | Mutation scheme |  |  |  |  |  |
| --- | --- | --- | --- | --- | --- | --- |
|  | Rieux [14] |  |  | Översti [13] |  |  |
|  | 50% | 95% | 99% | 50% | 95% | 99% |
| 1.2M growth | 387 | 3,835 | 7,361 | 295 | 2,869 | 5,603 |
| 1.2M const. | 177 | 761 | 1,148 | 152 | 661 | 1,006 |
| 300K const. | 193 | 859 | 1,293 | 149 | 675 | 1,085 |

These distributions are altered by conditioning on an observation of *m* matches in a randomly-sampled database of size *n* (Fig. 2, coloured curves). For the largest database we now see a clear difference between the two constant-size populations. For example *m* = 10 represents 0.1% of the database, consistent with 300 matches in the smaller population, a value that is well supported by the unconditional distribution and so the conditional distribution is centred around 300. However, 0.1% of the larger population is 1,200, which is not supported by the unconditional distribution and so the conditional distribution is shifted towards lower values, with most support between about 600 and 1,200. There is a similar effect for the *m* = 10 conditional distribution in the 1.2M growth population (note the different x-axis scale).

Estimated quantiles for the solid curves in the middle column of Fig. 2 are given in Table 2. For the other two demographic scenarios under the Översti mutation scheme [13], see Table A1 (300K constant) and Table A2 (1.2M growth). Corresponding quantiles for the Rieux mutation scheme [14] are given in Table A3 (1.2M constant), Table A4 (300K constant) and Table A5 (1.2M growth).

**Table 2.** Estimated quantiles of the number of matching individuals under the mutation scheme of [13]. Distributions shown in Fig. 2, middle column. *m* denotes the observed count of the haplotype in a database of size *n*. See text for references to additional tables for the other demographic scenarios.

| Quantile | 50% | 95% | 99% |
| --- | --- | --- | --- |
| Unconditional | 152 | 661 | 1,006 |
| $n = 100 / m = 0$ | 150 | 649 | 989 |
| $n = 1,000 / m = 0$ | 129 | 559 | 852 |
| $n = 10,000 / m = 0$ | 54 | 233 | 357 |
| $n = 100 / m = 1$ | 361 | 1,016 | 1,487 |
| $n = 1,000 / m = 1$ | 312 | 878 | 1,255 |
| $n = 10,000 / m = 1$ | 130 | 367 | 514 |
| $n = 100 / m = 2$ | 581 | 1,414 | 1,727 |
| $n = 1,000 / m = 2$ | 497 | 1,181 | 1,580 |
| $n = 10,000 / m = 2$ | 208 | 487 | 655 |
| $n = 1,000 / m = 5$ | 1,058 | 1,751 | 1,853 |
| $n = 10,000 / m = 5$ | 439 | 813 | 1,007 |
| $n = 10,000 / m = 10$ | 820 | 1,353 | 1,625 |

The number of meioses separating individuals with matching mitogenomes ranges up to a few hundred, and is almost never *>* 500 (Fig. 3). This is close to unrelated for most practical purposes, but random pairs of individuals are very unlikely to be related within 1,000 meioses, and so pairs with matching mitogenomes are much more closely related than average pairs of individuals. Key quantiles for the distributions of matching pairs are given in Table 3. As a guide for comparison, a coalescent theory approximation [18] for the mean numbers of meioses separating a random pair are 100K and 400K for our small and large constant-size populations, respectively.

**Figure 3.**
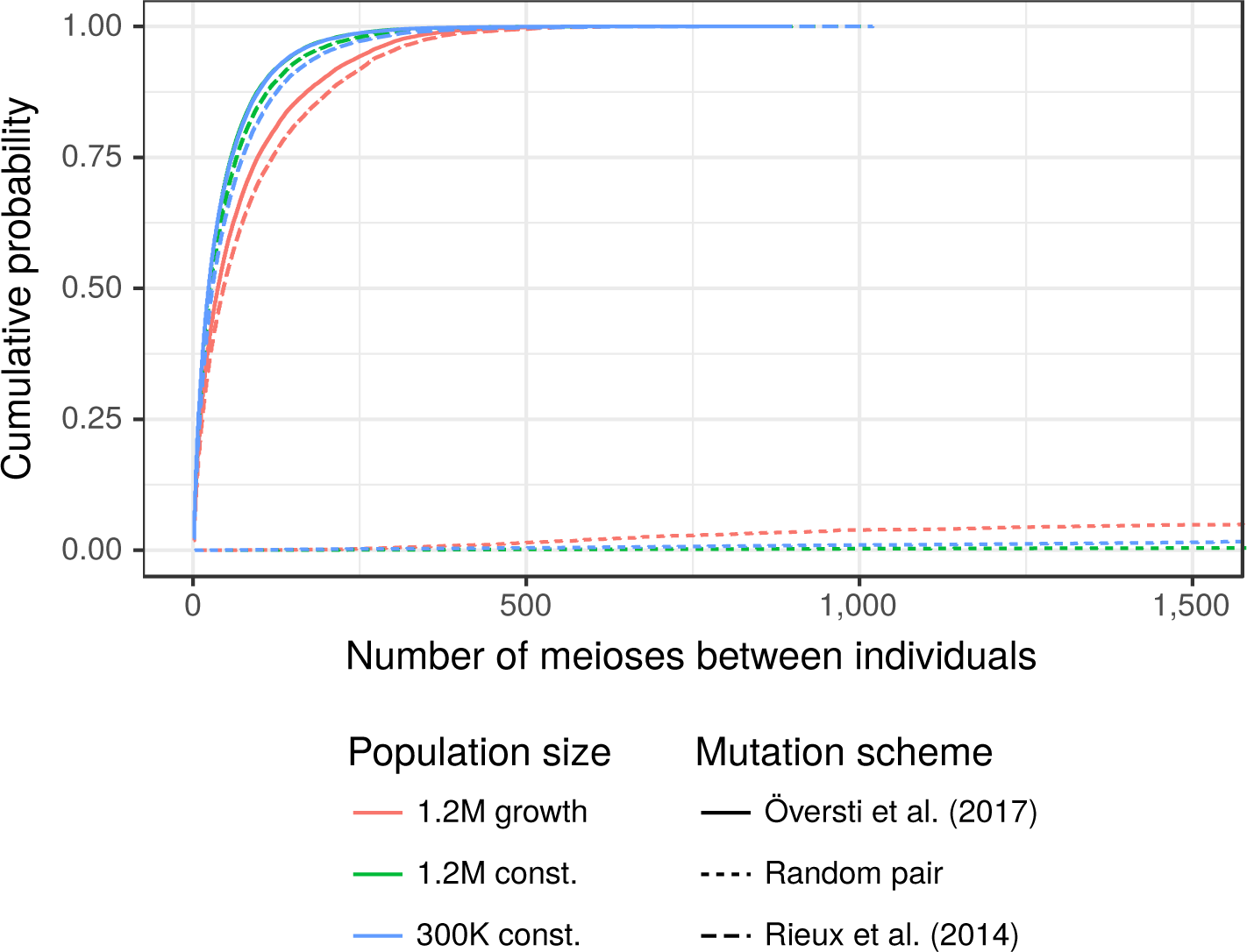
Number of meioses between pairs of individuals. The dotted lines correspond to random pairs of individuals, the solid and dashed lines are for pairs observed to have matching mitogenomes. See Table 3 for quantiles.

**Table 3.** Estimated quantiles of the number of meioses between pairs of individuals with matching mitogenome. Quantiles of the distributions shown in Fig. 3 (solid and dashed curves).

| Demographic scenario | Mutation scheme |  |  |  |  |  |
| --- | --- | --- | --- | --- | --- | --- |
|  | Rieux [14] |  |  | Översti [13] |  |  |
|  | 50% | 95% | 99% | 50% | 95% | 99% |
| 1.2M growth | 46 | 294 | 434 | 37 | 262 | 377 |
| 1.2M const. | 27 | 177 | 304 | 23 | 155 | 266 |
| 300K const. | 29 | 198 | 341 | 23 | 154 | 272 |

## Discussion

Empirical mitogenome databases do not in practice represent random samples from a well-defined population, so that detailed comparisons with our simulation models are not meaningful. However, we have verified here that the haplotype diversity generated by our simulation models is broadly comparable with that observed in two real databases from large populations.

In our related paper on Y profile matching [11], we showed that because of the high mutation rates of contemporary Y profiles, the numbers of males with Y profile matching a PoI (person of interest) are low, typically up to a few tens, and that this number is little affected by population size or growth. Moreover the clusters of matching males are related within a few tens of meioses and so are unlikely to be randomly distributed in the population relevant to a typical crime scene. We argued that it was therefore not appropriate to report a match probability (a special case of the likelihood ratio) to measure the weight of evidence, even though likelihood ratios are central to the evaluation of autosomal DNA profiles.

In the present paper we have shown that the situation for mtDNA evidence is intermediate between Y and autosomal profiles. Because the whole-mitogenome mutation rate is an order of magnitude smaller than the mutation rate for contemporary Y profiles, the number of individuals matching a PoI is correspondingly larger, and varies more with demography. The unconditional distribution (Table 1) is very similar for the two constant-size populations that differ in size by a factor of four, but for the growing population the median number of matches is about twice as big. As for the case of Y profiles, our simulation-based approach can easily take into account information from a frequency database, although this requires the assumption that the database is a random sample from the population, which is rarely the case in practice.

The mitolina software that we have presented here can be used to inform the evaluation of the weight of mtDNA evidence in forensic applications, similar to our recommended approach to presenting Y-profile evidence: simulation models are used to obtain a conservative estimate of the number of individuals sharing the evidence sample mitogenome, with conditioning on a database frequency if available. Current methods for evaluating mtDNA evidence rely directly on a database count of the observed mitogenome [3], and are affected by poor representativeness of the databases, and its limited informativeness when there are many rare mitotypes. Our approach can also make use of a database count of the haplotype, but this information is used to adjust an unconditional distribution and so is less sensitive to the database size and sampling scheme.

Limitations of our analysis include the range of demographic scenarios that we can consider, and the difficulty in assessing which demographic scenario is appropriate for any specific crime. Our assumption of neutrality is unlikely to be strictly accurate [19], nor our assumption of a generation time of 25 years, constant over generations. We used two mutation rate schemes [13, 14] based on phylogenetic estimates, as no pedigree-based mutation rates were available for the entire mitogenome. Some discrepancy has been noted between the two estimation methods [20], and the rate may have changed over time [21]. If contemporary pedigree-based mutation rates become available we could improve our mutation model, but that would not address mutation rate changes over time. We have not here addressed the case of mixed mtDNA samples or heteroplasmy (multiple mitogenomes arising from the same individual).

While we have focussed our examples on human populations because of the important role of the mitogenome in human identification and relatedness testing, with appropriate modifications of the demographic model, mitolina and the methods described here can be used for non-human applications of mtDNA. Examples include tracking the source of ivory [22], other areas of wildlife forensics [23] and inferences about the demographic histories of natural populations [24].

## Materials and Methods

### Mitogenome mutation models

We simulated the mitogenome as a binary sequence subject to neutral mutations, using the rates estimated by both Rieux et al. (2014) [14] and Översti et al. (2017) [13], shown in Table 4. They both partitioned the mitogenome into four regions: hypervariable 1+2 (HVS1 + HVS2), protein coding codon 1+2 (PC1 + PC2), protein coding codon 3 (PC3), and ribosomal-RNA + transfer-RNA (rRNA + tRNA). However, the HVS1 + HVS2 region of [14] consisted of 698 sites whereas that of [13] had 1,122 sites, although their total mutation rate estimates for the region are similar.

**Table 4.** Mutation rates per site and per 10^7^ generations. *L* and *U* denote lower and upper bounds of a 95% highest posterior density interval. The values here are 25 times the per-year rates of [14, 13], because we assume 25-year generations

| Region | Rieux et al. 2014 [14] |  | Översti et al. 2017 [13] |  |
| --- | --- | --- | --- | --- |
| | # sites | ($L, U$) | # sites | ($L, U$) |
| HVS1 + HVS2 | 698 | (56.40, 100.76) | 1,122 | (31.23, 72.53) |
| PC1 + PC2 | 7,565 | (1.43, 2.34) | 7,565 | (2.92, 6.00) |
| PC3 | 3,776 | (6.42, 10.19) | 3,776 | (4.80, 10.53) |
| rRNA + tRNA | 4,031 | (1.89, 3.17) | 4,031 | (2.35, 5.75) |
| Mitogenome | 16,070 | (2.16, 11.64) | 16,494 | (2.40, 13.84) |

### Population simulations

We simulated populations of mitogenomes under three demographic scenarios. Two constant-size Wright-Fisher populations, of 50K and 200K females per generation, were simulated for 1,200 generations. The third scenario started with a constant female population size of 10,257 for 1,000 generations, followed by growth at a rate at 2% per generation over 150 generations to reach a final generation with 200K females. Following [11], individuals in the final three generations are considered to be “live”, and in those generations males were also simulated making total live population sizes of 300K, 1.2M and 1.2M. All the females in any generation had the same distribution of offspring number (no between-female variation in reproductive success).

We assigned mitogenomes to the founders randomly with replacement from a US Caucasian database of 263 mitogenomes (259 distinct haplotypes, see Fig. 1) [15], coding each site as 0 if it matched the rCRS reference sequence [8], and 1 otherwise. Each mother-child transmission was subject to mutation, which changed a 0 to a 1, and vice versa. The same mutation rate was assigned to each site within each region, sampled from a normal distribution with 95% interval from Table 4.

The mean whole-mitogenome mutation rate per generation was 0.0135 for [13] and 0.0110 for [14], or about 1 mutation per 74 generations and 1 per 90 generations, respectively. Therefore, following one line of descent over 1,200 generations, the expected numbers of mutations to affect the mitogenome are 16.3 using [13] and 13.2 using [14]. The probabilities that there is any site affected by two mutations and so reverts to its original state during those 1,200 generations are 0.024 and 0.033, respectively.

We simulated five population under each of the three demographic scenarios. For each population simulation and both mutation models, we conducted five replicates of the sequence evolution process: assigning sequences to the founders and then mutations at each meiosis. Thus, for each mutation model and demographic scenario, 25 live populations of mitogenomes were created. In each live population, a PoI (person of interest) was randomly drawn 10,000 times, and we recorded how many live individuals had the same mitogenome as the PoI. Thus, a total of 5 × 5× 10K = 250K PoIs were sampled for each mutation and demography combination. Further, for 10% of the PoI, the number of meioses between the PoI and each matching individual was recorded.

Following the methodology of [11], in addition to the unconditional distribution of the number of mitogenome matches between a PoI and another live individual, we use importance sampling reweighting to approximate the distribution conditional on observing the PoI mitogenome *m* times in a database of size *n*, assumed to have been chosen randomly in the population.

Software to perform these simulations is implemented in the open-source R packages `mitolina` [25, 26], based on Rcpp [27], and `malan` [28], previously used for Y profile simulations [11].

## Acknowledgements

We thank Walther Parson, Adrien Rieux, Sanne Översti, Charla Marshall, Kimberly Andreaggi, and Miroslava Derenko for helpful responses to our queries. This work was supported in part by the Otto Mønsted Foundation and a short term fellowship from the International Society for Forensic Genetics (ISFG).

## Supplementary Material

**Table A1:**
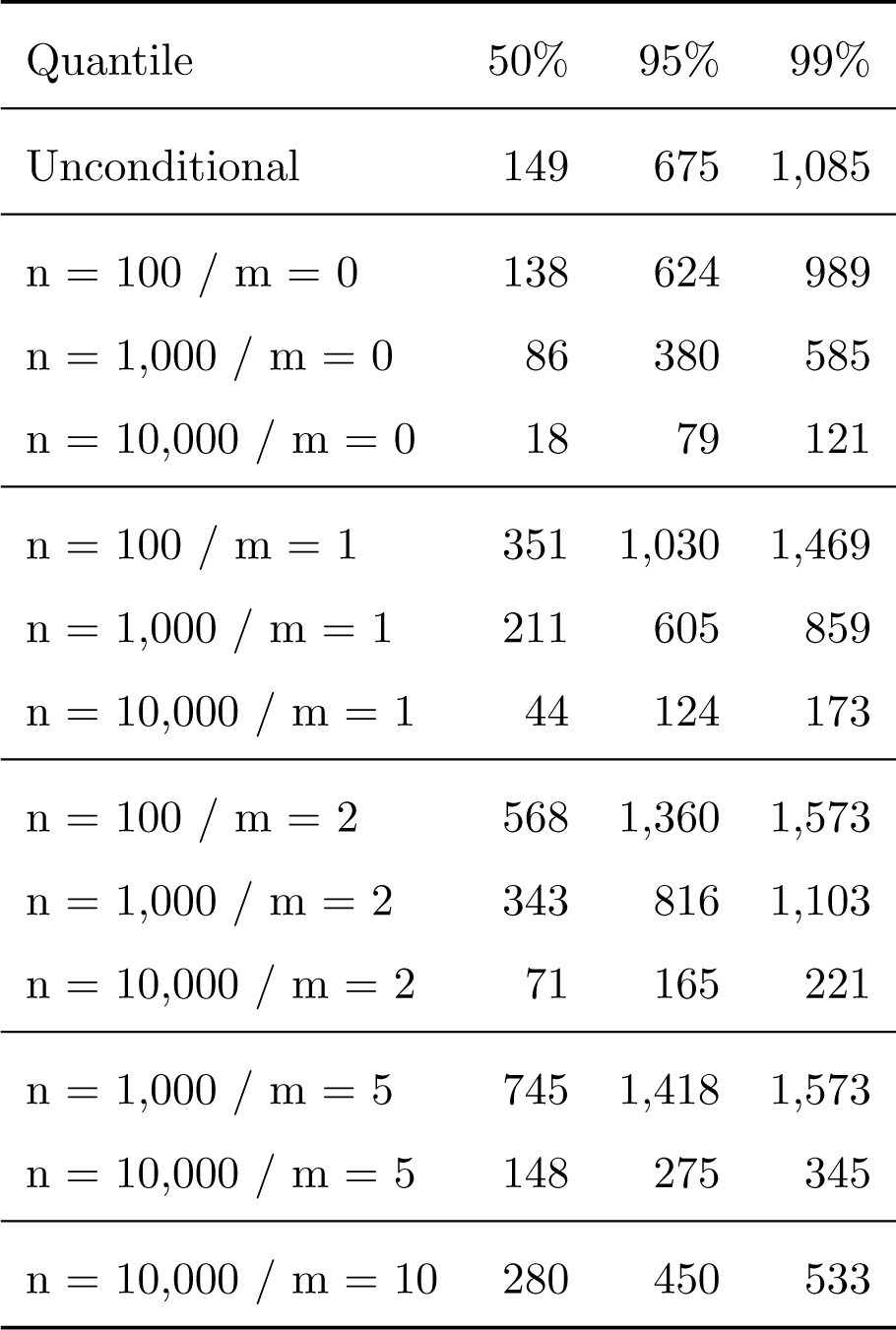
Approximate quantiles of the number of matching individuals. Key quantiles of the distributions shown in Fig. 2 for the mutation scheme of Översti [13], and for the 300K constant demographic scenario.

**Table A2:** Approximate quantiles of the number of matching individuals. Key quantiles of the distributions shown in Fig. 2 for the mutation scheme of Översti [13], and for the 1.2M growth demographic scenario.

| Quantile | 50% | 95% | 99% |
| --- | --- | --- | --- |
| Unconditional | 295 | 2,869 | 5,603 |
| n = 100 / m = 0 | 268 | 2,524 | 4,655 |
| n = 1,000 / m = 0 | 161 | 1,134 | 2,126 |
| n = 10,000 / m = 0 | 46 | 231 | 375 |
| n = 100 / m = 1 | 1,548 | 6,042 | 9,108 |
| n = 1,000 / m = 1 | 661 | 2,556 | 3,665 |
| n = 10,000 / m = 1 | 130 | 406 | 588 |
| n = 100 / m = 2 | 3,246 | 9,108 | 10,561 |
| n = 1,000 / m = 2 | 1,372 | 3,683 | 5,340 |
| n = 10,000 / m = 2 | 223 | 569 | 782 |
| n = 1,000 / m = 5 | 3,567 | 7,168 | 9,177 |
| n = 10,000 / m = 5 | 534 | 1,038 | 1,302 |
| n = 10,000 / m = 10 | 1,084 | 1,762 | 2,140 |

**Table A3:** Approximate quantiles of the number of matching individuals. Key quantiles of the distributions shown in Fig. 2 for the mutation scheme of Rieux [14], and for the 1.2M constant demographic scenario.

| Quantile | 50% | 95% | 99% |
| --- | --- | --- | --- |
| Unconditional | 177 | 761 | 1,148 |
| n = 100 / m = 0 | 174 | 744 | 1,114 |
| n = 1,000 / m = 0 | 146 | 627 | 956 |
| n = 10,000 / m = 0 | 56 | 244 | 375 |
| n = 100 / m = 1 | 416 | 1,154 | 1,627 |
| n = 1,000 / m = 1 | 352 | 981 | 1,364 |
| n = 10,000 / m = 1 | 137 | 386 | 543 |
| n = 100 / m = 2 | 658 | 1,528 | 2,136 |
| n = 1,000 / m = 2 | 558 | 1,297 | 1,725 |
| n = 10,000 / m = 2 | 219 | 514 | 686 |
| n = 1,000 / m = 5 | 1,154 | 2,151 | 2,293 |
| n = 10,000 / m = 5 | 463 | 856 | 1,061 |
| n = 10,000 / m = 10 | 862 | 1,364 | 1,639 |

**Table A4:** Approximate quantiles of the number of matching individuals. Key quantiles of the distributions shown in Fig. 2 for the mutation scheme of Rieux [14], and for the 300K constant demographic scenario.

| Quantile | 50% | 95% | 99% |
| --- | --- | --- | --- |
| Unconditional | 193 | 859 | 1,293 |
| n = 100 / m = 0 | 176 | 784 | 1,190 |
| n = 1,000 / m = 0 | 99 | 432 | 676 |
| n = 10,000 / m = 0 | 18 | 81 | 124 |
| n = 100 / m = 1 | 440 | 1,222 | 1,605 |
| n = 1,000 / m = 1 | 242 | 702 | 982 |
| n = 10,000 / m = 1 | 45 | 128 | 179 |
| n = 100 / m = 2 | 704 | 1,517 | 1,827 |
| n = 1,000 / m = 2 | 391 | 932 | 1,228 |
| n = 10,000 / m = 2 | 73 | 169 | 226 |
| n = 1,000 / m = 5 | 836 | 1,507 | 1,818 |
| n = 10,000 / m = 5 | 151 | 285 | 355 |
| n = 10,000 / m = 10 | 290 | 458 | 545 |

**Table A5:** Approximate quantiles of the number of matching individuals. Key quantiles of the distributions shown in Fig. 2 for the mutation scheme of Rieux [14], and for the 1.2M growth demographic scenario.

| Quantile | 50% | 95% | 99% |
| --- | --- | --- | --- |
| Unconditional | 387 | 3,835 | 7,361 |
| n = 100 / m = 0 | 339 | 3,242 | 5,662 |
| n = 1,000 / m = 0 | 182 | 1,291 | 2,342 |
| n = 10,000 / m = 0 | 47 | 237 | 386 |
| n = 100 / m = 1 | 2,004 | 7,697 | 11,463 |
| n = 1,000 / m = 1 | 756 | 2,875 | 4,164 |
| n = 10,000 / m = 1 | 133 | 415 | 608 |
| n = 100 / m = 2 | 4,027 | 11,275 | 14,221 |
| n = 1,000 / m = 2 | 1,544 | 4,133 | 5,579 |
| n = 10,000 / m = 2 | 228 | 586 | 806 |
| n = 1,000 / m = 5 | 3,926 | 7,799 | 9,608 |
| n = 10,000 / m = 5 | 552 | 1,057 | 1,332 |
| n = 10,000 / m = 10 | 1,095 | 1,779 | 2,134 |

**Figure A1.**
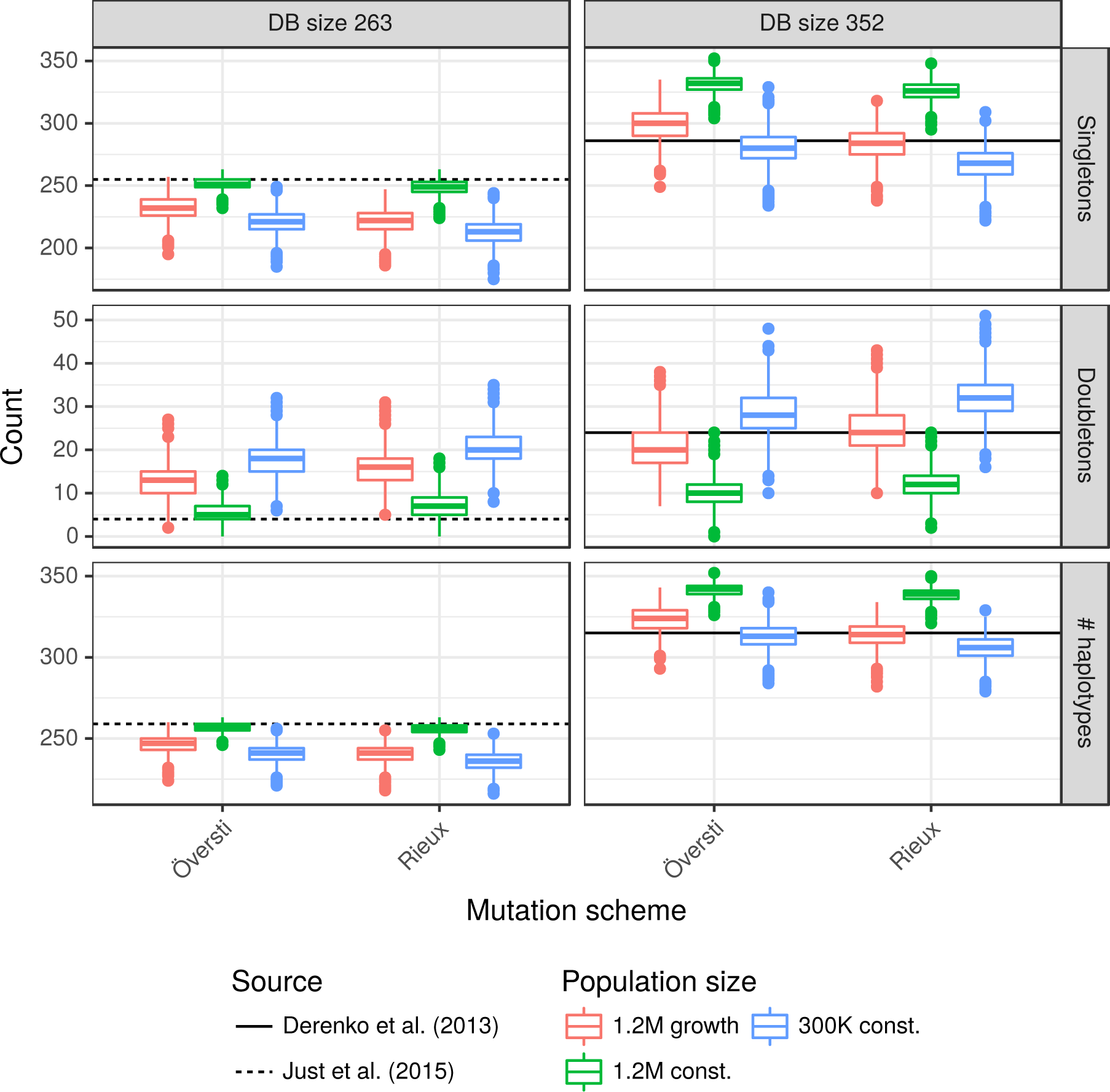
Comparison of simulated with US and Iranian databases. The distribution of the numbers of singletons, doubletons and distinct haplotypes in 2,500 random databases of sizes 263 and 351 obtained under our three demographic and two mutation models. The horizontal reference lines are from [15, 16]. [16] does not provide number of singletons and doubletons, but these numbers (286 and 24, respectively) were obtained directly from the authors.

## References

[1] John M Butler and Barbara C Levin. Forensic applications of mitochondrial DNA. Trends in Biotechnology, 16(4):158–162, 1998.

[2] A Carracedo, W Bär, P Lincoln, W Mayr, N Morling, B Olaisen, P Schneider, B Budowle, B Brinkmann, P Gill, M Holland, G Tully, and M Wilson. DNA Commission of the International Society for Forensic Genetics: guidelines for mitochondrial DNA typing. Forensic Science International, 110(2):79–85, 2000.

[3] W. Parson, L. Gusmão, D.R. Hares, J.A. Irwin, W.R. Mayr, N. Morling, E. Pokorak, M. Prinz, A. Salas, P.M. Schneider, and T.J. Parsons. DNA Commission of the International Society for Forensic Genetics: Revised and extended guidelines for mitochondrial DNA typing. Forensic Science International: Genetics, 13:134–142, 2014.

[4] M. Thomas P. Gilbert, Toomas Kivisild, Bjarne Grønnow, Pernille K. Andersen, Ene Metspalu, Maere Reidla, Erika Tamm, Erik Axelsson, Anders Götherström, Paula F. Campos, Morten Rasmussen, Mait Metspalu, Thomas F. G. Higham, Jean-Luc Schwenninger, Roger Nathan, Cees-Jan De Hoog, Anders Koch, Lone Nukaaraq Møller, Claus Andreasen, Morten Meldgaard, Richard Villems, Christian Bendixen, and Eske Willerslev. Paleo-eskimo mtdna genome reveals matrilineal discontinuity in greenland. Science, 320(5884):1787–1789, 2008.

[5] Tim H. Heupink, Sankar Subramanian, Joanne L. Wright, Phillip Endicott, Michael Carrington Westaway, Leon Huynen, Walther Parson, Craig D. Millar, Eske Willerslev, and David M. Lambert. Ancient mtdna sequences from the first australians revisited. Proceedings of the National Academy of Sciences, 113(25):6892–6897, 2016.

[6] Jennifer D. Churchill, Dixie Peters, Christina Capt, Christina Strobl, Walther Parson, and Bruce Budowle. Working towards implementation of whole genome mitochondrial DNA sequencing into routine casework. Forensic Science International: Genetics Supplement Series, 6:e388–e389, 2017.

[7] Christina Strobl, Mayra Eduardoff, Magdalena M. Bus, Marie Allen, and Walther Parson. Evaluation of the precision ID whole mtDNA genome panel for forensic analyses. Forensic Science International: Genetics, 35:21–25, 2018.

[8] Richard M Andrews, Iwona Kubacka, Patrick F Chinnery, Robert N Lightowlers, Douglass M Turnbull, and Neil Howell. Reanalysis and revision of the Cambridge reference sequence for human mitochondrial DNA. Nature Genetics, 23, 1999.

[9] C.D. Steele and D. Balding. Weight of evidence for forensic DNA profiles. Wiley, 2nd edition, 2015.

[10] T King, G Fortes, P Balaresque, M Thomas, D Balding, P Delser, R Neumann, W Parson, M Knapp, S Walsh, L Tonasso, J Holt, M Kayser, J Appleby, P Forster, D Ekserdjian, M Hofreiter, and K Schörer. Identification of the remains of King Richard III. Nat. Commun., 5:5631, 2014.

[11] Mikkel Meyer Andersen and David J Balding. How convincing is a matching Y-chromosome profile? PLOS Genetics, 13(11):e1007028, 2017.

[12] S Willuweit and L Roewer. The New Y Chromosome Haplotype Reference Database. Forensic Science International: Genetics, 15:43–48, 2015.

[13] Sanni Översti, Päivi Onkamo, Monika Stoljarova, Bruce Budowle, Antti Sajantila, and Jukka U. Palo. Identification and analysis of mtDNA genomes attributed to Finns reveal long-stagnant demographic trends obscured in the total diversity. Scientific Reports, 7, 2017.

[14] Adrien Rieux, Anders Eriksson, Mingkun Li, Benjamin Sobkowiak, Lucy A. Weinert, Vera Warmuth, Andres Ruiz-Linares, Andrea Manica, and François Balloux. Improved Calibration of the Human Mitochondrial Clock Using Ancient Genomes. Molecular Biology and Evolution, 31(10):2780–2792, 2014.

[15] Rebecca S. Just, Melissa K. Scheible, Spence A. Fast, Kimberly Sturk-Andreaggi, Alexander W. Röck, Jocelyn M. Bush, Jennifer L. Higginbotham, Michelle A. Peck, Joseph D. Ring, Gabriela E. Huber, Catarina Xavier, Christina Strobl, Elizabeth A. Lyons, Toni M. Diegoli, Martin Bodner, Liane Fendt, Petra Kralj, Simone Nagl, Daniela Niederwieser, Bettina Zimmermann, Walther Parson, and Jodi A. Irwin. Full mtGenome reference data: Development and characterization of 588 forensic-quality haplotypes representing three U.S. populations. Forensic Science International: Genetics, 14:141–155, 2015.

[16] Miroslava Derenko, Boris Malyarchuk, Ardeshir Bahmanimehr, Galina Denisova, Maria Perkova, Shirin Farjadian, and Levon Yepiskoposyan. Complete mitochondrial dna diversity in iranians. PLOS ONE, 8(11):1–14, 11 2013.

[17] Ellen D. Gunnarsdöttir, Mingkun Li, Marc Bauchet, Knut Finstermeier, and Mark Stoneking. High-throughput sequencing of complete human mtDNA genomes from the Philippines. Genome Res., 21(1):1–11, 2011.

[18] John Wakeley. Coalescent Theory: An Introduction. Roberts & Co, 2008.

[19] J. William O. Ballard and David M. Rand. The population biology of mitochondrial DNA and its phylogenetic implications. Annual Review of Ecology, Evolution, and Systematics, 36(1):621–642, 2005.

[20] Neil Howell, Christy Bogolin Smejkal, D.A. Mackey, P.F. Chinnery, D.M. Turnbull, and Corinna Herrnstadt. The pedigree rate of sequence divergence in the human mitochondrial genome: There is a difference between phylogenetic and pedigree rates. The American Journal of Human Genetics, 72(3):659–670, 2003.

[21] Brenna M. Henn, Christopher R. Gignoux, Marcus W. Feldman, and Joanna L. Mountain. Characterizing the time dependency of human mitochondrial DNA mutation rate estimates. Molecular Biology and Evolution, 26(1):217–230, 2009.

[22] I Yasuko, N Georgiadis, H Tomoko, and A. Roca. Triangulating the provenance of African elephants using mitochondrial DNA. Evolutionary Applications, 6(2):253–265, 2012.

[23] A Linacre. Application of mitochondrial dna technologies in wildlife investigation - species identification. Forensic Sci Rev, 18(1):1–8, 2006.

[24] L. A. Rollins, A. P. Woolnough, R. Sinclair, N. J. Mooney, and W. B. Sherwin. Mitochondrial DNA offers unique insights into invasion history of the common starling. Molecular Ecology, 20(11):2307–2317, 2011.

[25] R Development Core Team. R: A Language and Environment for Statistical Computing. R Foundation for Statistical Computing, Vienna, Austria, 2018. ISBN 3-900051-07-0.

[26] Mikkel Meyer Andersen. mitolina: Mitochondrial lineage analysis. https://github.com/mikldk/mitolina, 2018.

[27] D Eddelbuettel and JJ Balamuta. Extending *R* with *C++*: A Brief Introduction to *Rcpp*. PeerJ Preprints, 5:e3188v1, aug 2017.

[28] Mikkel M Andersen. malan: MAle Lineage ANalysis. The Journal of Open Source Software, 3(25), 2018.

